# Hot days are associated with short-term adrenocortical responses in a Southern African arid-zone passerine bird

**DOI:** 10.1101/2021.02.17.431578

**Authors:** Lesedi L. Moagi, Amanda R. Bourne, Susan J. Cunningham, Ray Jansen, Celiwe A. Ngcamphalala, Amanda R. Ridley, Andrew E. McKechnie

## Abstract

Non-invasive methods for investigating the biological effects of environmental variables are invaluable for understanding potential impacts of climate change on behavioural and physiological stress responses of free-ranging animals. Foraging efficiency, body mass maintenance and breeding success are compromised in Southern pied babblers *Turdoides bicolor* exposed to air temperatures between ^~^35°C and ^~^40°C. We tested the hypothesis that these very hot days are acute stressors for free-ranging babblers by quantifying the relationship between daily maximum air temperature (T_max_) and faecal glucocorticoid metabolite (fGCM) levels. We collected naturally-excreted droppings from free-ranging pied babblers and analysed fGCM levels. Levels of fGCMs in droppings collected after 3pm were independent of same-day T_max_ < 38 °C and averaged 140.25 ng g^−1^ Dry Weight ± 56.92 ng g^−1^ DW (mean ± SD) over this range. Above an inflection T_max_ = 38 °C, however, fGCM levels increased linearly with same-day T_max_ and averaged 190.79 ng g^−1^ DW ± 70.13 ng g^−1^ DW. There was no relationship between T_max_ on the previous day and fGCM levels in droppings collected the following morning. Group size, breeding stage, sex and rank did not predict variation in fGCM levels. These results suggest that very high T_max_ may be linked to acute, but not chronic, heat stress responses. The fGCM levels we measured are likely to represent a biologically meaningful response to an environmental stressor (high T_max_), suggesting a physiological mechanism underlying observed changes in behaviour and reproductive success at high temperatures in this species.

## Introduction

Stress responses mediated by the endocrine system are a vital component of animals’ reaction to environmental perturbations (Sheriff, Dantzer, Delehanty, Palme, & Boonstra. 2011). Responses triggered by exposure to a stressor also include physiological and behavioural changes (Agarwal & Prabhakaran, 2005; Stott, 1981), which contribute to an animal’s ability to respond appropriately to environmental change (Jensen, Moseby, Paton, & Fanson, 2019) and its likelihood of survival (Asres & Amha, 2014). Environmental temperature is an important determinant of stress in animals (de Bruijn & Romero, 2011; Jessop et al., 2016; Krause et al., 2016; Xie et al., 2017; Ruuskanen 2019), and understanding species-specific stress responses to increasingly frequent and severe heat waves is important for effective conservation and management under advancing climate change.

Glucocorticoids are a class of steroid hormones important for energy mobilisation, immune function, and metabolism and are a crucial part of stress responses (MacDougall-Shackleton, Bonier, Romero, & Moore, 2019). There are two primary glucocorticoids: i) cortisol, found in ungulates, carnivores and primates, and ii) corticosterone, found in rodents, birds and reptiles (Palme et al 2005; Touma & Palme 2005). Quantification of these glucocorticoids is an established and popular method that has been used to study stress responses in animals in relation to external stressors (Ganswindt et al., 2012; Crino et al., 2016; Dantzer et al., 2010 & Sheriff et al., 2011), including high temperatures (Xie et al., 2017).

Stress responses are usually categorised as either acute or chronic. Acute stress is associated with the rapid, transient release of glucocorticoids, often to levels far above baseline, in response to a specific stimulus (Beiko et al., 2004). These short-term elevations can be beneficial, supporting immune responses and mobilising energy reserves which may, for example, allow an individual to escape a potentially dangerous situation such as an attempted predation (Buchanan, 2000). Chronic stress, on the other hand, occurs over longer periods and involves prolonged elevation of glucocorticoid levels in response to ongoing exposure to stressors. Negative effects of chronic stress include disrupted cognition, immune function, and reproduction, which potentially have long term consequences such as reduced breeding success (McEwen, 2004; Dhabhar, 2009 & Lupien et al., 2009).

Levels of circulating glucocorticoids have often been determined directly using blood samples (Crino et al., 2020; MacDougall-Shackleton et al., 2019). However, blood collection can be stressful for the study animal (Pavlova et al., 2018; Small et al., 2017), and handling stress risks artificially elevating measurements of stress hormones. Alternative non-invasive methods based on quantifying glucocorticoid metabolites in droppings, hair, feathers, or saliva have attracted much attention and have been validated for an increasing number of species (Sheriff et al., 2011; Palme, 2019; Dantzer et al., 2010; Hämäläinen, Heistermann, Fenosoa & Kraus, 2014), including Southern pied babblers (*Turdoides bicolor*; Jepsen et al, 2019). Measurement of adrenocortical activity via faecal glucocorticoid metabolites (fGCMs) can eliminate the need for handling of study animals altogether, thereby avoiding artificially increasing the circulating glucocorticoid levels of the study animals (Hodges et al. 2010), provided that the collection of droppings is not stressful to the animal. In some study populations, collecting droppings also presents an opportunity to sample more frequently and under natural conditions (Jepsen et al., 2019). Non-invasive measurements of stress responses have important conservation applications because they can be used to investigate the sub-lethal effects of social and ecological stressors under natural conditions (Wikelski & Cooke, 2006; Sherif et al., 2011; Narayan, 2013 & Dantzer et al., 2014). Although increasing in popularity among researchers, non-invasive sampling of glucocorticoids in avian droppings (especially from free-ranging birds) have been used less frequently than in studies of mammals and reptiles (Palme, 2019).

Little is known about avian stress responses to high air temperatures. We know that chronic exposure to heat can have strongly negative effects on birds. These effects include a trade-off between shade-seeking, resting, heat dissipation behaviour and foraging, leading to reduced foraging effort, efficiency and success (du Plessis et al. 2012, Edwards et al 2015, Cunningham et al 2015, van de Ven et al 2019), body mass loss at high temperatures (du Plessis et al 2012, Sharpe et al 2019, van de Ven 2019), reduced breeding success (Salaberria et al 2014, Wiley & Ridley 2016, DuRant et al 2019, Sharpe et al 2019, van de Ven et al 2020, Bourne et al 2020a), reduced survival of adults (Bourne et al 2020c, Sharpe et al 2019), mass mortality events (McKechnie & Wolf 2010 & Conradie et al 2020) and population declines (Iknayan & Beissinger 2018). Acute exposure to heat can also result in elevated metabolic rates, dehydration, and death (McKechnie 2019). A study on Sonoran Desert birds showed that plasma corticosterone was not elevated during summer compared to winter, even during a year when summer temperatures were higher than normal (Wingfield et al., 1992). However, an Australian study found elevated corticosterone levels after exposure to heat in some species but not others (Xie et al 2017).

In this study, we test the relationship between temperature and fGCM levels in free-ranging Southern pied babblers (*Turdoides bicolor*). Specifically, we test the hypothesis that high maximum daily air temperatures (T_max_), associated with compromised foraging and net 24-hr mass loss (du Plessis *et al*., 2012) as well as failed reproduction (Bourne et al., 2020a) in pied babblers, will trigger an acute stress response in this species. Pied babblers are known to lose mass overnight and reduce provisioning to nestlings when T_max_ ≥ 35.5°C (du Plessis et al 2012, Wiley & Ridley 2016) and are unable to breed successfully at average T_max_ ≥ 38°C (Bourne et al., 2020a). We therefore tested a specific hypothesis that temperatures in the mid-to high-30s (°C) would trigger an acute stress response in this species. We expected fGCM levels to increase during hot weather. In order to elucidate the time scale over which fGCM levels were elevated in response to hot weather and the extent to which stress responses to an extremely hot day are acute or chronic, we compared the effect of T_max_ on fGCM levels in samples collected that afternoon, and the effect of T_max_ of the preceding day (T_maxPrev_) on fGCM levels in samples collected in the morning. We reasoned that, if an extremely hot day triggers an acute stress response, T_max_ would influence pied babblers’ fGCM levels in the afternoon of the same day but not in the morning on the following day. Alternately, if an extremely hot day triggers a chronic stress response with carry-over effects to the subsequent day, then elevated fGCM levels would remain detectable the following day.

## Methods and Materials

### Study site

The study was conducted at Kuruman River Reserve, located in the southern Kalahari Desert, 28 km west of Van Zylsrus town in the Northern Cape Province, South Africa (S 26° 58′ E 21° 49″). The study site’s annual average summer maximum temperature from 2005 to 2019 was 34.2 ± 0.9 °C (range: 32.4– 36.5 °C) and summer rainfall averaged 185.4 ± 86.2 mm (range: 64.4–352.1 mm; Bourne et al., 2020b). The 33-km^2^ reserve is flat with a sandy substrate supporting semi-arid savannah vegetation (Mucina and Rutherford, 2006).

### Study species

Pied babblers are medium-sized (60 – 90g) cooperatively breeding passerines (Ridley 2016, Bourne et al., 2019). As part of an on-going study of the behaviour of wild pied babblers habituated to the presence of human observers, the birds were observed under natural conditions (Ridley, 2016). During our study, pied babbler group sizes ranged from 2 – 8 adults. Adults are defined as individuals aged ≥ 12 months (Raihani & Ridley, 2007). Habituation of the pied babblers in the study population allows human observation from distances of 1 – 5m (Ridley & Raihani, 2007), enabling collection of droppings (Bourne et al., 2019). The birds are ringed with metal and coloured rings for individual identification, which makes it easy to assign each faecal sample to an individual. Each group has a dominant female and male, with the remaining individuals being subordinates (Nelson-Flower et al., 2011). Pied babblers have high reproductive skew, with 95% of young produced by the dominant pair (Nelson-Flower et al., 2011). The pied babblers are highly vocal and primarily terrestrial foragers that inhabit open woodlands; these are characteristics that make them easy to observe for research purposes (Ridley, 2016).

### Data collection

Droppings were collected by following the birds and sampling excreta within 1 min after defecation by a known individual bird, with the droppings immediately transferred to a screw-cap, plastic Eppendorf tube sealed with parafilm (following Bourne et al., 2019). The date and time of collection of the sample, and the identity of each bird and group were recorded (following Jepson et al., 2019). Each babbler group was visited weekly during the breeding season and monitoring visits lasted up to 90 min; which was sufficient for proper identification of individuals and collection of the droppings. Faecal samples were collected throughout the day during one austral summer breeding season from November 2018 to February 2019 (n = 898 samples in total). Samples were frozen at −18°C within 1.81 ± 1.07 h of collection (mean ± SD hours, range = 0.02 – 11.38 h). Samples were collected from approximately 71 individual pied babblers from 18 groups, including dominant and subordinate adults of both sexes. Weather data were obtained from a weather station onsite (Vantage Pro2, Davis Instruments, Hayward, U.S.A.), which recorded air temperature (Ta; °C), wind speed (m s^−1^), rainfall (mm) and solar radiation (W m^−2^) at 10-min intervals throughout the study period (van de Ven, McKechnie & Cunningham, 2019).

### Faecal glucocorticoid metabolites analysis

Of the 898 samples collected, we selected a subset of 228 for analysis. We selected all of the samples collected after 3 pm (afternoon samples, n = 114) and then selected another 114 from the samples collected before 9 am (morning samples). Using afternoon samples collected after 3pm allowed for sufficient time between exposure to a temperature stressor during the hottest time of the day and detecting a measurable response in fGCM levels in babblers after ^~^ 2h (see Jepsen et al., 2019). Afternoon samples were collected on days distributed across a range of T_max_ (28 °C – 41 °C), and from individuals from different group sizes (2-8 adults), sexes (n = 43 males, 63 females, 8 unknown sex), ranks (n = 52 dominant, 62 subordinate) breeding stages (n = 45 from breeding groups, 69 from non-breeding groups). In order to identify whether any stress response detected represented an acute or chronic response, we further analysed 114 morning samples across the same temperature range of 28 °C – 41 °C, recorded on the previous day (T_maxPrev_), and similarly distributed across group sizes (3-8 adults), sex (n = 58 males, 56 females), rank (n = 52 dominant, 62 subordinate), and breeding stage (n = 45 breeding, 69 non-breeding). Morning samples were randomly selected within the categories breeding stage, group size, sex and rank. Individuals from pairs (group size = 2 adults) were excluded from statistical analysis because the samples were too few (n= 1). Morning and afternoon samples were not paired.

Frozen faecal samples were lyophilized, pulverized and sieved to remove undigested material before adding 1.5 ml of 80% ethanol in distilled water to 0.050–0.055 g of faecal powder and vortexing for 15 min to facilitate steroid extraction (Ganswindt et al., 2002). After centrifuging for 10 min, the samples were transferred into microcentrifuge tubes and stored at −20 °C (Jepsen et al., 2019). Immunoreactive fGCMs were quantified using an enzyme immunoassay (EIA) utilizing an antibody against 5ß-pregnane-3,11ß,21-triol-20-one-CMO:BSA (tetrahydrocorticosterone). Characteristics of the EIA including cross-reactivities are given by Quillfeldt and Möstl (2003). This EIA was validated for the reliable quantification of fGCMs in pied babblers by Jepsen et al. (2019).

The coefficients of variation (CV) for the inter-assay variance of the subset of 32 afternoon samples ranged from 10.17% to 10.36% and the CV for intra-assay ranged from 6.33% to 6.64%, while the remaining 82 afternoon samples had CV for inter-assay that were 9.03% to 11.17% and the CV for intra-assay variance ranged from 6.33% to 6.64%. The 114 morning samples CV for the inter-assay variance were 12.5% and 14.5% while CV for intra-assay variance were 5.30% and 7.75%. The combined CV for inter-assay variance for all 228 samples were 13.26% and 13.62%. The sensitivity of the enzyme immune-assay used for all 228 samples is 9 ng/g faecal dry weight.

## Statistical analyses

We used R version 3.6.3 for all analyses (R Core Team, 2020). We checked model residuals of all the variables for normality using QQ plots. The variance inflation factor (VIF) was used to assess collinearity between numeric variables (Harrison et al., 2018) and all VIF ≤ 2. Collinearity between categorical and numeric predictor variables was tested using analysis of variance (ANOVA). In the subset of the afternoon samples, sex and group size were associated (F1,99 = 5.920, *p* = 0.016), with more subordinate females than males in the group sizes ranging from 4 to 8 individuals. Group size and breeding stage were associated in the morning samples (F_2,100_ = 4.830, *p* = 0.009), with larger groups more likely to be breeding (also see Ridley 2016, Bourne et al. 2020b). Correlation between two categorical predictors was checked using a Chi-Square test and no pairs of categorical variables were correlated. Correlated predictors were not included in the same additive models (Harrison et al., 2018) and T_max_ and T_maxPrev_ were not correlated.

We tested the effects of T_max_ (for afternoon samples), T_maxPrev_ (for morning and afternoon samples), breeding stage, group size, rank, sex and the interaction between rank and sex on fGCM levels in linear mixed-effects models (LMMs) fitted using the lme4 package (Bates et al., 2015b, Harrison et al., 2018). We included group identity included as a random term. The inclusion of individual identity as a random terms in addition to group identity resulted in unstable models and of the two random terms, group identity explained the greatest proportion of variation while avoiding destabilising the models (Grueber, Nakagawa, Laws, & Jamieson, 2011; Harrison et al., 2018). Effects of both T_max_ and T_maxPrev_ were tested in an additive model for afternoon samples in order to explore the relative importance of acute (same day temperature) vs chronic (temperature on previous day) responses. The interaction between rank and sex was included to confirm the association found by Jepsen et al (2019), whereby dominant male pied babblers exhibited higher fGCM levels, using a larger dataset collected in a different year. Data are presented as mean estimate ± 1 standard error (SE) unless otherwise stated. The three continuous explanatory variables were rescaled by centring and standardising by the mean of the variables for model comparisons (Harrison et al., 2018). Model terms with confidence intervals not intersecting zero were considered to explain significant patterns in our data (Grueber et al., 2011). Models were compared using Akaike’s Information Criterion (AIC) with the *MuMIn* package and competing models within ΔAICc 2.0 of the top model were averaged (Bartoń, 2016, Harrison et al., 2018). Model-averaged coefficients are presented following van de Ven et al., (2020) & Harrison et al., (2018).

When visual inspection of the data suggested a non-linear response, we supplemented the above LMM analyses with a segmented linear regression using the R package *segmented* (Muggeo, 2008). Segmented regression can be used to identify temperature thresholds (‘breakpoints’) above which fGCM levels begin to increase. We analysed the effect of T_max_ separately above and below the identified breakpoints, including the random term for group.

## Results

For afternoon samples, T_max_ significantly predicted fGCM levels, with higher fGCM levels at higher maximum temperatures (Table 1, Figure 1). The single best-fit model explaining variation in fGCM levels had a model weight of 0.779 (Table 1) and included both T_max_ and T_maxPrev_. We identified a breakpoint at T_max_ = 38.0 °C (Figure 1). There was no effect of T_max_ at temperatures < 38.0 °C but fGCM levels increased with increasing T_max_ at temperatures > 38.0 °C. We found no significant effect of group size, breeding stage, rank, sex or the interaction between rank and sex on afternoon fGCM levels (Table 1). For morning samples, there was no significant relationship between fGCM levels and any of the potential predictors (Table 2). Dominant females had higher fGCM levels on average than the other birds but the trend was not statistically significant.

**Figure 1:**
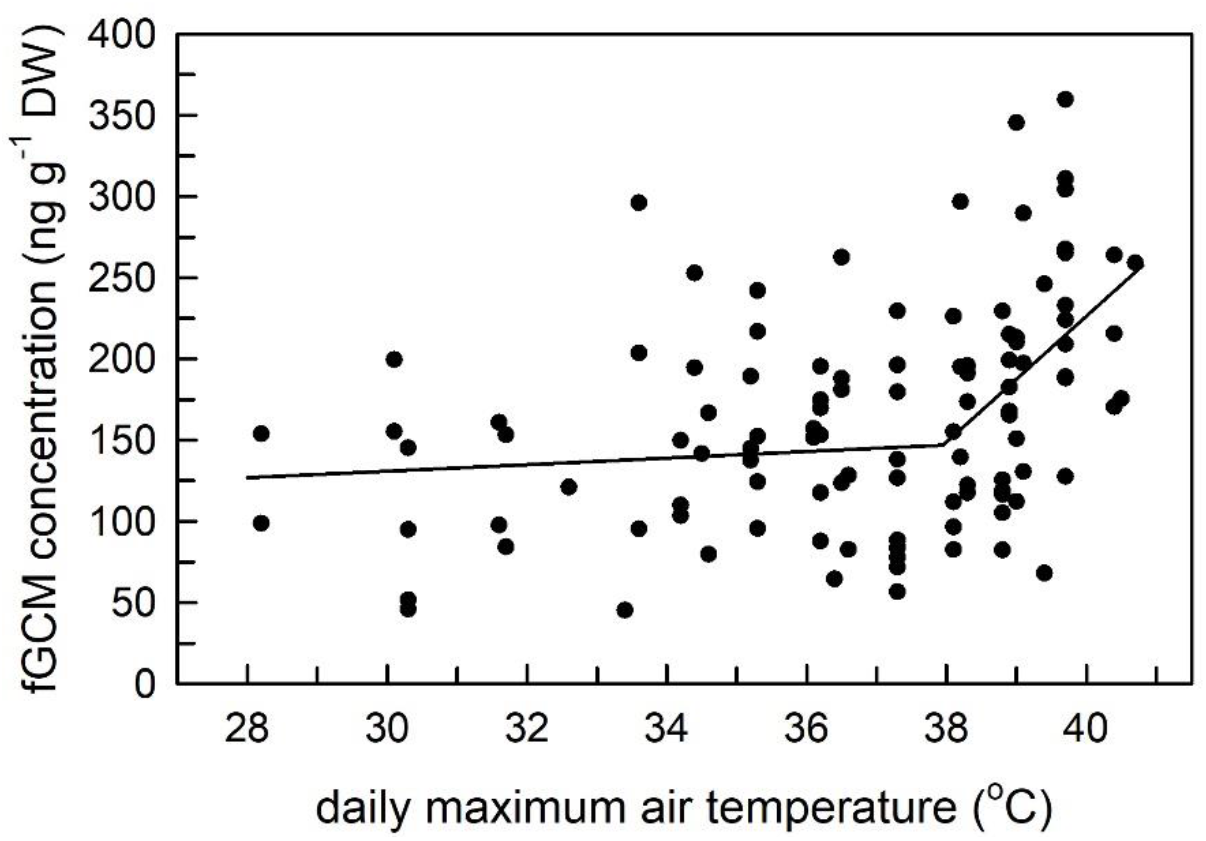
Relationship between afternoon fGCM levels and T_max_ with increasing fGCM levels as T_max_ increases in Southern Pied Babblers (Turdoides bicolor), with a non-linear effect of T_max_ on fGCM levels. The break point has been identified at T_max_ = 38.0 °C (95% CI:36.876, 39.122). Data represented was taken from 144 samples collected from 71 individual Southern Pied Babblers from 18 groups, including dominant and subordinate adults of both sexes.

**Table 1:** Model outputs for analyses of fGCM levels for afternoon samples. Data comes from 101 afternoon samples from 48 different individuals in 15 groups. Estimate, standard error and confidence intervals are shown for the top model. Significant terms are shown in bold. Null models shown for comparison with top model sets. Random terms: 1 | Group.

| Models | AICc | $\Delta AICc$ | $\omega i$ |
| --- | --- | --- | --- |
| $T_{max} + T_{maxPrev}$ | 1110.1 | 0.00 | 0.779 |
| $T_{max}$ | 1114.3 | 4.17 | 0.097 |
| Breeding Stage | 1114.4 | 4.30 | 0.091 |
| Rank*Sex | 1116.5 | 6.35 | 0.033 |
| $T_{maxPrev}$ | 1123.6 | 13.48 | 0.001 |
| Sex | 1128.0 | 17.91 | 0.000 |
| Group Size | 1128.3 | 18.20 | 0.000 |
| Rank | 1128.5 | 18.38 | 0.000 |
| Null model | 1133.6 | 23.52 | 0.000 |
| Effect size of explanatory terms in the top model | Estimate | SE | 95% CI |
| Intercept | 165.266 | 9.554 | 146.349/184.618 |
| $T_{max}$ | **28.197** | **8.975** | **10.592/45.629** |
| $T_{maxPrev}$ | -3.715 | 8.906 | -21.055/13.657 |

**Table 2:** Model outputs for analyses of fGCM levels for morning samples. Data comes from 103 morning samples from 56 different individuals in 18 groups, collected before 9am. Estimate, standard error and confidence intervals are shown for the top model. Significant terms are shown in bold. Null models shown for comparison with top model sets. Random terms: 1 | Group.

| Models | AICc | $\Delta AICc$ | $\omega i$ |
| --- | --- | --- | --- |
| Rank * Sex | 1043.4 | 0.00 | 0.990 |
| Rank | 1054.6 | 11.23 | 0.004 |
| Sex | 1055.6 | 12.29 | 0.002 |
| Breeding Stage | 1056.1 | 12.79 | 0.002 |
| T<sub>maxPrev</sub> | 1056.4 | 13.00 | 0.001 |
| Group Size | 1157.6 | 14.21 | 0.001 |
| Null model | 1060.2 | 16.81 | 0.000 |
| Effect size of explanatory terms in the top model | Estimate | SE | 95% CI |
| Intercept | 199.532 | 9.228 | 101.633/137.431 |
| Rank (rank = SUB) | -21.200 | 11.836 | -44.158/1.757 |
| Sex (sex = MALE) | -18.110 | 12.175 | -41.725/5.506 |
| Rank*Sex | 18.423 | 16.472 | -13.525/50.371 |

We followed procedures similar to those described in Jepsen et al (2019) and we found no relationship between the amount of time between collection and freezing and measured fGCM levels for either morning (F1.101 = 1.127; *p* = 0.312) or afternoon samples (F1.107= 0.087, *p* = 0.769).

## Discussion

Our results reveal that high daily maximum air temperatures are associated with elevated fGCM levels and may be linked to acute heat stress in pied babblers. Specifically, we found a significant effect of T_max_ on fGCM levels of samples collected that afternoon, and no significant effect of T_maxPrev_ on fGCM levels levels of samples collected either in the morning or the afternoon of the following day, suggesting that high T_max_ triggers an acute stress response in the pied babblers. We found no relationship between fGCM levels and group size or breeding stage and, unlike the Jepsen et al (2019) study, we did not detect a significant relationship between fGCM levels and rank or sex, or the interaction between the two.

A significant increase in fGCM levels was evident at T_max_ ≥ 38°C. This T_max_ is similar to several thresholds related to the effects of T_max_ on body mass and breeding success in the same population of pied babblers. Du Plessis et al. (2012) found that T_max_ ≥ 35.5 °C has the potential to compromise the ability of pied babblers to maintain body condition. Pied babblers typically lose ^~^4% of their body mass overnight, but due to reduced foraging success on hot days they may fail to gain at least that amount during the day on days ≥ 35.5 °C and this can result in gradual deterioration of body condition over time (du Plessis et al., 2012). Wiley & Ridley (2016) reported reduced provisioning to pied babbler nestlings at temperatures ≥ 35.5 °C and Bourne et al (2020a) recently reported, using 15 years of life history data, that no pied babbler young survived when mean T_max_ during the nestling period exceeded 38°C. Together, these observational studies suggest that pied babblers may experience heat stress when daily T_max_ exceeds 35.5 °C and approaches 40 °C. Our finding, that fGCM levels increase sharply in free-living pied babblers under natural conditions when T_max_ exceeds 38°C, is consistent with these observed temperature thresholds, and suggests a physiological mechanism.

Evidence for the existence of critical air temperatures above which costs are incurred is present in several other bird species. For example, Edwards et al. (2015) found that temperature had a significant effect on the behaviour of Western Australian Magpie *Cracticus tibicen dorsalis*, with temperatures exceeding 27 °C resulting in a significant decline in foraging effort. Sharpe et al. (2019) found that Jacky Winters *Microeca fascinans* (a small Australian passerine), were never observed foraging, provisioning nestlings or even incubating, at air temperatures > 38 °C (also see Bayter, 2019). Similarly, van de Ven et al. (2019) showed that foraging efficiency in Yellow-billed Hornbills was negatively affected by both panting and increasing time spent in cooler microsites, resulting in lower numbers of prey captured overall at high air temperature despite no change in foraging effort. The threshold temperature above which breeding male hornbills did not gain any mass over a 12 h day was 38.4 °C. In common fiscals *Lanius collaris*, increasing numbers of days of T_max_ > 37 °C negatively influenced fledgling tarsus length and increasing number of days of T_max_ > 35 °C were associated with nestlings having to stay in the nest longer before they were ready to fledge (Cunningham et al., 2013). The presence of threshold temperatures similar to those of pied babblers in other species suggests that these other species might also be experiencing acute physiological stress detectable in fGCM levels.

Although our data were all collected within a single summer, we were able to collect samples across a range of daily maximum temperatures and from a large number of different individuals. In addition, by sampling individuals varying in sex, rank, breeding stage and group size, we were able to evaluate potential causes of individual variation in fGCM levels, to differentiate this from temperature effects. A potential limitation of our study is that delays between collecting and freezing the samples could affect measurements of fGCM levels. Exposure to biotic and abiotic elements has influenced measured fCGM levels in some other studies (Lafferty et al., 2019). Measured fGCM levels may also be influenced by environmental factors, for example ambient temperature and humidity between excretion and freezing of the sample can affect bacterial metabolisation that can result in possible increases or decreases in the fGCM levels (Terio et al., 2002, Washburn & Millspaugh, 2002, Palme et al., 2013). Differences in microclimates across the range of a mammal species, the American pika *Ochotona princeps*, influenced fGCM levels when droppings were not immediately collected and preserved after defecation (Wilkening, 2016). Despite these risks, we did not find any significant influence of the period between collection and freezing on fGCM levels in this study (also see Jepsen et al., 2019).

The use of a non-invasive sampling technique with a free-living study population avoids the potential confounds associated with the influence of capture, handling, and captivity on fGCM levels (Wingfield et al., 1995; Dickens et al., 2009). By collecting droppings from the ground after they were naturally excreted by the bird and after the bird had moved away on its own (Bourne et al, 2019; Jepsen et al. 2019), we entirely avoided handling the study animals and eliminated the potentially confounding effect of capture stress (Harper & Austad 2000, Millspaugh 2001 & Touma & Palme 2005). Jepsen et al.’s (2019) study compared captive and free-ranging wild pied babblers and found that wild individuals had significantly lower baseline fGCM levels, even after captive birds had been habituated to captivity for four days. Our measured fGCM levels, collected from free-living pied babblers, are likely to represent a biologically meaningful response to an environmental stressor – in this case high T_max_ – experienced under natural conditions.

Non-invasive physiological methods can be very useful for behavioural ecologists who wish to explore physiological correlates of behavioural strategies without disrupting the behaviour of their study organisms to collect these data (Bourne et al., 2019). FGCM levels can be correlated directly with observed natural behaviour and environmental variables. Wider application of non-invasive techniques could open new avenues for assessing behavioural and physiological responses concurrently in wild animals under natural conditions, an increasingly important capability as ecologists seek to understand the impacts of global change pressures on local animal populations (Stillman, 2019). Measured fGCM levels in droppings are less affected by episodic fluctuations of hormone secretion than those in blood and might, therefore, represent the animal’s hormonal status more accurately than blood samples (Touma et al., 2004). Droppings are also a particularly useful matrix for measuring fGCM levels because they are relatively abundant and can be collected with minimal disturbance to study animals (Möstl & Palme, 2002, Millspaugh, & Washburn, 2004 and Palme, 2019). FGCM levels in faecal samples can be a reliable indicator of the levels present in blood (Palme, 2019) and birds make particularly useful study subjects as most bird species excrete both urine and droppings together – effectively resulting in a double peak of hormone levels.

The avian upper lethal body temperature is 45 - 46°C, just 4 – 5 °C above resting body temperature (Dawson & Schmidt – Nielsen 1964, Prinzinger et al., 1991). Our study showed that non-invasive methods through faecal sampling can help us understand the physiological impacts of high temperatures on avian species. We demonstrated that fGCM levels in free-living pied babblers, are elevated above a threshold maximum daily air temperature of 38 °C, a result that is not confounded with or influenced by handling or captivity stress. This threshold temperature is very similar to previously published threshold temperatures for pied babblers which demonstrate that high temperatures can be detrimental for reproductive success and survival (Bourne et al 2020a, 2020c). Behavioural adjustments and compromised survival may, therefore, be partially explained by an acute physiological stress response to high temperatures. As anthropogenic climate change is advancing rapidly (IPCC, 2014), studying the biological effects of climate change on free-ranging animals using non-invasive methods can be helpful in terms of understanding the potential impact on their behavioural and physiological responses.

## Acknowledgements

We thank Camilla Soravia and Sabrina Engesser for helping with the collection of pied babbler droppings during 2018-2019. Abongile Ndzungu, Stefanie Ganswindt and Nicole Hagenah were instrumental in performing the EIAs in the Endocrine Research Laboratory at University of Pretoria. We thank the management teams at the Kuruman River Reserve (KRR) and surrounding farms, Van Zylsrus, South Africa, for making the work possible. The KRR was financed by the Universities of Cambridge and Zurich, the MAVA Foundation, and the European Research Council (Grant No. 294494 to Tim Clutton-Brock), and received logistical support from the Mammal Research Institute of the University of Pretoria. This work was funded by the DST-NRF Centre of Excellence at the FitzPatrick Institute for African Ornithology, the University of Cape Town, the Oppenheimer Memorial Trust (Grant No. 20747/01 to ARB), the British Ornithologists’ Union, the Australian Research Council (Grant No. FT110100188 to ARR), and the National Research Foundation of South Africa (Grant No. 110506 to SJC & Grant No. 119754 to LLM). Any opinions, findings and conclusions or recommendations expressed in this material are those of the authors and do not necessarily reflect the views of the National Research Foundation.

## References

Asres, A. & Amha, N. (2014). Effects of stress on animal health: A Review. Journal of Biology. 4(27): 2224–3208.

Agarwal, A. & Prabhakaran S.A. (2005). Mechanism, measurement and prevention of oxidative stress in male reproductive physiology. Indian Journal of Experimental Biology. 43: 963–974.

Bates, D., Maechler, M., Bolker, B. & Walker, S. (2015b). Fitting linear mixed-effects models using lme4. Journal of Statistics Software. 67: 1–48.

Bartoń, K. (2016). MuMIn: multi-model inference. R package Version 1.15.6. Available at https://CRAN.R-project.org/package=MuMIn.

Bayter, C. (2019). Effects of ambient temperature on incubation behaviour and parental body condition in jacky winters (*Microeca fascinans*). – Unpublished Honours dissertation, Australian National University.

Beiko, J., Lander, R., Hampson, E., Boon, F. & Cain, D.P. (2004). Contribution of sex differences in the acute stress response to sex differences in water maze performance in the rat. Behavioural Brain Research, 151(1-2): 239–253.

Bourne, A.R., McKechnie A.E., Cunningham S.J., Ridley A.R., Woodborne S.M. & Karasov W.H. (2019). Non-invasive measurement of metabolic rates in wild, free living birds using doubly labelled water. Functional Ecolology. 33: 162–174. doi: 10.1111/1365-2435.13230.

Bourne, A.R., Cunningham, S.J., Spottiswoode, C.N. & Ridley, A.R. (2020a). High temperatures drive offspring mortality in a cooperatively breeding bird. Proceedings of the Royal Society B: Biological Sciences 287(1931): 20201140 doi.org/10.1098/rspb.2020.1140.

Bourne, A.R., Cunningham, S.J., Spottiswoode, C.N. & Ridley, A.R. (2020b). Compensatory Breeding in Years Following Drought in a Desert-Dwelling Cooperative Breeder. Frontiers in Ecology and Evolution 8: 190. doi:10.3389/fevo.2020.00190.

Bourne, A.R., Cunningham, S.J., Spottiswoode, C.N. & Ridley, A.R. (2020c). Hot droughts compromise interannual survival across all group sizes in a cooperatively breeding bird. Ecology Letters. doi.org/10.1111/ele.13604.

Boyles, J. G., Seebacher, F., Smit, B. & McKechnie, A. E. (2011). Adaptive thermoregulation in endotherms may alter responses to climate change. – Integrated Comparative Biology. 51: 676–690.

Buchanan, K.L. (2000). Reply from KL Buchanan. Trends in Ecology & Evolution, 15(10): 419.

Clutton-Brock, T.H., MacColl, A.D.C., Chadwick, P., Gaynor, D., Kansky, R. & Skinner, J. D. (1999). Reproduction and survival of suricates, *Suricata suricatta*, in the southern Kalahari. African Journal of Ecology, 37: 69 e80.

Conradie, S.R., Woodborne, S.M., Cunningham, S.J. & McKechnie, A.E., (2019). Chronic, sublethal effects of high temperatures will cause severe declines in southern African arid-zone birds during the 21st century. Functional Ecology. 116(28): 14065–14070 doi.org/10.1073/pnas.1821312116.

Conradie, S.R., Woodborne, S.M., Wolf, B.O., Pessato, A., Mariette, M.M. & McKechnie, A.E. (n.d). (2020). Avian mortality risk during heat waves will increase greatly in arid Australia during the 21^st^ Century. Conservation Physiology. 8, coaa04.

Crino, O.L., Buchanan, K.L., Trompf, L., Mainwaring, M.C. & Griffith, S.C. (2016). Stress reactivity, condition, and foraging behavior in zebra finches: effects on boldness, exploration, and sociality. General and Comparative Endocrinology. doi:10.1016/j.ygcen.2016.01.014.

Crino, O.L., Driscoll, S.C., Brandl, H.B., Buchanan, K.L. & Griffith, S.C. (2020). Under the weather: corticosterone levels in wild nestlings are associated with ambient temperature and wind. General and Comparative Endocrinology, 285: 113247. doi:10.1016/j.ygcen.2019.113247.

Cunningham, S.J., Martin, R.O., Hojem, C.L. & Hockey, P.A.R. (2013). Temperatures in Excess of Critical Thresholds Threaten Nestling Growth and Survival in a Rapidly-Warming Arid Savanna: A Study of Common Fiscals. PLoS ONE 8(9): e74613. doi:10.1371/journal.pone.0074613.

Cunningham, S.J., Martin, R.O. & Hockey, P.A.R. (2015). Can behaviour buffer the impacts of climate change on an arid-zone bird? Ostrich: Journal of African Ornithology 86(1&2): 119–126. doi.org/10.2989/00306525.2015.1016469.

Dantzer, B., McAdam, A.G., Palme, R., Fletcher, Q.E., Boutin, S., Humphries, M.M. & Boonstra, R. (2010). Fecal cortisol metabolite levels in free-ranging North American red squirrels: Assay validation and the effects of reproductive condition. General and Comparative Endocrinology, 167(2): 279–286. doi:10.1016/j.ygcen.2010.03.024.

Dawson W. R. (1954). Temperature regulation and water requirements of the brown and Abert towhees, *Pipilo fuscus* and *Pipilo aberti*. In University of California publications in zoology, Volume. 59 (eds. Bartholomew, G.A., Crescitelli, F., Bullock, T.H., Furgason, W.H. & Schechtman A.M.), pp. 81–123 Berkeley, CA: University of California Press.

Dawson, W.R. (1982). Evaporative losses of water by birds. Comparative Biochemistry Physiology A: Comparative Physiology. 71(4): 495–509.

Dawson, W.R. & Schmidt-Nielsen, K. (1964). Terrestrial animals in dry heat: desert birds. In: Dill DB (ed) Handbook of physiology: adaptation to the environment. American Physiological Society, Washington, D.C., pp 481–492.

de Bruijn, R. & Romero, L.M., (2011). Behavioral and physiological responses of wild-caught European starlings (Sturnus vulgaris) to a minor, rapid change in ambient temperature. Comparative Biochemical and Physiology a-Molecular & Integrative Physiology. 160: 260–266.

Dickens, M.J., Earle, K.A. & Romero, L.M., (2009). Initial transference of wild birds to captivity alters stress physiology. General Comparative Endocrinology. 160(1): 76–83.

du Plessis, K. L., Martin, R. O., Hockey, P. A. R., Cunningham, S. J. & Ridley, A. R. (2012). The costs of keeping cool in a warming world: Implications of high temperatures for foraging, thermoregulation and body condition of an arid-zone bird. Global Change Biology, 18(10): 3063–3070.

DuRant, S.E., Willson, J.D. & Carroll, R.B. (2019). Parental effects and climate change: will avian incubation behavior shield embryos from increasing environmental temperatures? Integrative Compartive Biology. 59: 1068–1080. doi:10.1093/icb/icz083.

Edwards, E.K., Mitchell, N.J. & Ridley, A.R. (2015). The impact of high temperatures on foraging behaviour and body condition in the Western Australian Magpie Cracticus tibicen dorsalis. 86(1-2): 137–144.

Ganswindt, A., Heistermann, M., Borragan, S. & Hodges, J.K. (2002). Assessment of testicular endocrine function in captive African elephants by measurement of urinary and fecal androgens. Zoo Biology 21(1): 27–36.

Grueber, C.E., Nakagawa, S. Laws, R.S. & Jamieson, I.G. (2011). Multimodal inference in ecology and evolution: challenges and solution. Journal of Evolution Biology. 24: 699–711. doi:10.1111/j.1420-9101.2010.02210.

Hämäläinen, A., Heistermann, M., Fenosoa, Z.S.E. & Kraus, C. (2014). Evaluating capture stress in wild gray mouse lemurs via repeated fecal sampling: method validation and the influence of prior experience and handling protocols on stress responses. General and Comparative Endocrinology, 195: 68–79. doi:10.1016/j.ygcen.2013.10.017.

Harrison, X.A., Donaldson, L., Co Correa-cano, M.E., Evans, J., Fisher, D.N., Goodwin, C.E.D., Robinson, B.S., Hodgson, D.J. & Inger, R. (2018). A brief introduction to mixed effects modelling and multi-model inference in ecology. Peer-reviewed journal, 6: 1–32.

Harper, J.M. & Austad, S.N. (2000). Fecal glucocorticoids: a noninvasive method of measuring adrenal activity in wild and captive rodents. Physiological and Biochemical Zoology 73: 12–22.

Hodges, K., Brown, J. & Heistermann, M. (2010). Endocrine monitoring of reproduction and stress. In: Wild mammals in captivity. (eds. Kleiman, D.G., Thompson, K.V. & Baer, C.K.) Chicago: University of Chicago Press. pp 447–468.

Iknayan, K.J. & Beissinger, S.R. (2018). Collapse of a desert bird community over the past century driven by climate change. – Proc. Natl Acad. Sci. USA 115: 8597–8602.

IPCC. (2013). Climate Change 2013: The Physical Science Basis. Contribution of Working Group I to the Fifth Assessment Report of the Intergovernmental Panel on Climate Change (eds. Stocker, T.F., D. Qin, G.-K. Plattner, M. Tignor, S.K. Allen, J. Boschung, A. Nauels, Y. Xia, V. Bex and P.M. Midgley). Cambridge University Press, Cambridge, United Kingdom and New York, NY, USA, 1535 pp. See https://www.ipcc.ch/report/ar5/wg1/

IPCC. (2014). Climate Change 2014: Synthesis Report. Contribution of Working Groups I, II and III to the Fifth Assessment Report of the Intergovernmental Panel on Climate Change. (eds C.W. Team, R.K. Pachauri & L.A. Meyer). Geneva, Switzerland, 151 pp. See https://www.ipcc.ch/site/assets/uploads/2018/05/SYR_AR5_FINAL_full_wcover.pdf

Jensen, M.A., Moseby, K.E., Paton, D.C. & Fanson, K.V. (2019). Non-invasive monitoring of adrenocortical physiology in a threatened Australian marsupial, the western quoll (*Dasyurus geoffroii*). Conservation Physiology, 7(1), coz069. doi:10.1093/conphys/coz069.

Jepsen, E.M., Ganswindt, A., Ngcamphalala, A., Bourne, A.R., Ridley, A.R. & Mckechnie, A.E. (2019). Non-invasive monitoring of physiological stress in an afrotropical arid-zone passerine bird, the Southern pied babbler. General and Comparative Endocrinology 276: 60–68.

Jessop, T.S., Lane, M.L., Teasdale, L., Stuart-Fox, D., Wilson, R.S., Careau, V. & Moore, I.T. (2016). Multiscale evaluation of thermal dependence in the glucocorticoid response of vertebrates. American Naturalist. 188: 342–356.

Journal of Statistical Software 67(1): 1–48. doi10.18637/jss.v067.i01.

Jury, M.R. (2013). Climate trends in southern Africa. South African Journal of Science. Volume 109 1/2, 11 pp. doi.org/10.1590/sajs.2013/980.

Krause, J.S., Pérez, J.H., Chmura, H.E., Sweet, S.K., Meddle, S.L., Hunt, K.E., Gough, L. Boelman, N. & Wingfield, J.C. (2016). The effect of extreme spring weather on body condition and stress physiology in Lapland longspurs and white-crowned sparrows breeding in the Arctic. General and Comparative Endocrinology. 237: 10–18.

Lafferty, D.J.R., Zimova, M., Clontz, L., Hackländer, K & Scott Mills, L.S. (2019). Noninvasive measures of physiological stress are confounded by exposure. Scientific Reports 9: 19170 doi.org/10.1038/s41598-019-55715-5.

Laver, P.N., Ganswindt, A., Ganswindt, S.B. & Alexander, K.A., (2012). Non-invasive monitoring of glucocorticoid metabolites in banded mongooses (*Mungos mungo*) in response to physiological and biological challenges. General and Comparative Endocrinology 179 (2): 178–183.

MacDougall-Shackleton, S.A., Bonier, F., Romero, L.M., & Moore, I.T. (2019). Glucocorticoids and “stress” are not synonymous. Integrative Organismal Biology, 1(1). doi:10.1093/iob/obz017.

McEwen, B.S. (2004). Protection and damage from acute and chronic stress: allostasis and allostatic overload and relevance to the pathophysiology of psychiatric disorders. Annals of the New York Academy of Sciences, 1032(1): 1–7.

McKechnie, A.E., Hockey, P.A. & Wolf, B.O. (2012). Feeling the heat: Australian landbirds and climate change. Emu Austrial Ornithology 112(2): 1–7.

McKechnie, A.E. (2019). “Physiological and morphological effects of climate change,” in Effects of Climate Change on Birds. (eds. Dunn, P.O & Moller, A.P) Oxford: Oxford University Press. pp. 120–133. doi: 10.1093/oso/9780198824268.003.0010.

McKechnie, A.E. & Wolf, B.O. (2019). The physiology of heat tolerance in small endotherms. Physiology 34: 302–313. doi.org/10.1152/physiol.00011.2019.

Millspaugh, J.J., Woods, R.J., Hunt, K.E., Raedeke, J., Brundige, G.C., Warshurn, B.E. & Wasser, S.K. (2001). Fecal glucocorticoid assays and the physiological stress response in Elk. Wildlife Society Bulletin, pp. 899–907. (1973-2006), Volume. 29, Issue. 3, pp. 899-907. See https://www.jstor.org/stable/3784417

Millspaugh, J.J. & Washburn, B.E. (2004). Use of fecal glucocorticoid metabolite measures in conservation biology research: considerations for application and interpretation. General and Comparative Endocrinology 138: 189–199.

Möstl, E. & Palme, R. (2002). Hormones as indicators of stress. Domestic Animal Endocrinology 23: 67–74.

Mucina, L. & Rutherford, M.C., (2006). The Vegetation of South Africa. South African National Biodiversity Institute, Lesotho and Swaziland.

Nelson-Flower, M.J., Hockey, P.A.R., O’Ryan, C., Raihani, N.J., du Plessisa, M.A. & Ridley, A. (2011). Monogamous dominant pairs monopolize reproduction in the cooperatively breeding pied babbler. Behavioural Ecology. 22: 559–565. doi:10.1093.

Palme, R., Rettenbacher, S., Touma, C., El-Bahr, S. & Möstl, E., (2005). Stress hormones in mammals and birds: comparative aspects regarding metabolism, excretion, and non-invasive measurement in faecal samples. Ann. N. Y. Academic Science. 1040(1): 162–171.

Palme, R., Touma, C., Arias, N., Dominchin, M.F. & Lepschy, M. (2013). Steroid extraction: get the best out of faecal samples. Wien Tierarztl Monatsschr 100: 238–246.

Palme, R., (2019). Non-invasive measurement of glucocorticoids: advances and problems. Physiological Behaviour. 199: 229–243.

Pavlova, E.V, Alekseeva, G.S., Erofeeva, M.N., Vasilieva, N.A., Tchabovsky, A.V, Naidenko, S.V, & Severtsov, A.N. (2018). The method matters: the effect of handling time on cortisol level and blood parameters in wild cats. Journal of Experimental Zoology Part A: Ecological and Integrative Physiology, 329(3): 112–119. doi:10.1002/jez.2191.

Prinzinger, R., Prebmar, A. & Schleucher, E. (1991). Body temperature in birds. Comparative Biochemistry and Physiology Part A: Physiology 99(4): 499–506. doi:10.1016/03009629(91)90122-S.

Quillfeldt, P. & Möstl, E. (2003). Resource allocation in Wilson’s storm-petrels Oceanites oceanicus determined by measurement of glucocorticoid excretion. Acta Ethology 5: 115–122.

R Core Team. 2020 R: a language and environment for statistical computing. Vienna, Austria: R Foundation for Statistical Computing. See http://www.r-project.org.

Raihani, N.J. & Ridley, A.R. (2007). Adult vocalizations during provisioning: offspring response and post fledging benefits in wild pied babblers. Animal Behaviour, 74(5): 1303–1309 doi:10.1016/j.anbehav.2007.02.025.

Ridley, A.R. & Raihani, N.J. (2007a). Variable postfledging care in a cooperative bird: causes and consequences. Behavioral Ecology, 18: 994–1000.

Ridley, A.R. & Raihani, N.J. (2007). Facultative response to a kleptoparasite by the cooperatively breeding pied babbler. Behavioural Ecology. doi:10.1093/beheco/arl092.

Ridley, A.R. (2016). Southern Pied Babblers: the dynamics of cooperation and conflict in a group-living society. In: Cooperative Breeding: Studies of Ecology. (eds. Koenig, W.D. & Dickinson, J.S.). Evolution and Behaviour. Cambridge University Press, pp. 115–132.

Ruuskanen, S., Hsu, B.Y. & Nord, A. (2019). Endocrinology of thermoregulation in birds in a changing climate. General and Comparative Endocrinology. 10.32942/osf.io/jzam3.

Salaberria, C., Celis, P., López-Rull, I. & Gil, D. (2014). Effects of temperature and nest heat exposure on nestling growth, dehydration and survival in a Mediterranean hole-nesting passerine. Ibis (Lond. 1859) 156: 265–275. doi:10.1111/ibi.12121.

Sharpe, L., Cale, B. & Gardne, J.L. (2019). Weighing the cost: the impact of serial heatwaves on body mass in a small Australian passerine. Journal of Avian Biology doi: 10.1111/jav.02355

Sheriff, M.J., Dantzer, B., Delehanty, B., Palme, R. & Boonstra, R. (2011). Measuring stress in wildlife: techniques for quantifying glucocorticoids. Physiological Ecology. Oecologia 166: 869–887.

Small, T.W., Bebus, S.E., Bridge, E.S., Elderbrock, E.K., Ferguson, S.M., Jones, B.C., & Schoech, S.J. (2017). Stress-responsiveness influences baseline glucocorticoid levels: revisiting the under 3 min sampling rule. General and Comparative Endocrinology, 247: 152–165. doi:10.1016/j.ygcen.2017.01.028.

Steenkamp, C.J., Vogel, J.C., Fuls, A., van Rooyen, N., & van Rooyen, M.W. (2008). Age determination of Acacia erioloba trees in the Kalahari. Journal of Arid Environments, 72(4): 302–313 doi.org/10.1016/j.jaridenv.2007.07.015.

Stott, G.H. (1981). What is animal stress and how is it measured. Journal of Animal Science, Volume 52. Issue 1. pp. 150–153. doi.org/10.2527/jas1981.521150x.

Terio, K.A., Brown, J.L., Moreland, R. & Munson, L. (2002). Comparison of different drying and storage methods on quantifiable levels of fecal steroids in the cheetah. Zoo Biology 21: 215–222.

Touma, C. & Palme, R. (2005). Measuring fecal glucocorticoid metabolites in mammals and birds: the importance of validation. Annals of the New York Academy of Sciences 1046: 54–74.

Touma, C., Palme, R. & Sachser, N. (2004). Analyzing corticosterone metabolites in fecal samples of mice: a noninvasive technique to monitor stress hormones. Hormones and Behavior 45:10–22.

van de Ven, T.M.F.N., McKechnie, A.E. & Cunningham, S.J. (2019). The costs of keeping cool: behavioural trade-offs between foraging and thermoregulation are associated with significant mass losses in an arid-zone bird. Oecologia, 191(1): 205–215. doi:10.1007/s00442-019-04486-x.

van de Ven, T.M.F.N., McKechnie, A.E., Er, S. & Cunningham, S.J. (2020). High temperatures are associated with substantial reductions in breeding success and offspring quality in an arid-zone bird. Oecologia 193: 225–235.

Washburn, B.E. & Millspaugh, J.J. (2002). Effects of simulated environmental conditions on glucocorticoid metabolite measurements in white-tailed deer feces. General and Comparative Endocrinology 127: 217–222.

Wiley, E.M. & Ridley, A.R. (2016). The effects of temperature on offspring provisioning in a cooperative breeder. Animal Behavior. 117: 187–195. doi:10.1016/j.anbehav.2016.05.009.

Wilkening, J.L., Ray, C. & Varner, J. (2016). When can we measure stress noninvasively? Postdeposition effects on a fecal stress metric confound a multiregional assessment. Ecology and Evolution 6: 502–513.

Wingfield, J.C., Vleck, C.M. & Moore, M.C. (1992). Seasonal changes of the adrenocortical response to stress in birds of the Sonoran Desert. The Journal of Experimental Zoology 264: 419–428.

Wingfield, J.C., O’Reilly, K.M. & Astheimer, L.B. (1995). Modulation of the adrenocortical responses to acute stress in arctic birds: a possible ecological basis. Animal Zoology. 35(3): 285–294.

Xie, S., Romero, L.M., Htut, Z.W., & McWhorter, T.J. (2017). Stress responses to heat exposure in three species of Australian desert birds. Physiological and Biochemical Zoology, 90(3): 348–358 doi:10.1086/690484.

Xie, S., Woolford, L. & McWorther, T.J. (2020). Organ histopathology and hematological changes associated with heat exposure in Australian desert birds. Journal of Avian Medicine and Surgery 34(1): 41–51.

